# Amino acid exchangeability and surface accessibility underpin the effects of single substitutions

**DOI:** 10.1101/2025.06.13.659595

**Authors:** Berk A. Alpay, Piyush Nanda, Emma Nagy, Michael M. Desai

**Affiliations:** Department of Systems Biology, Harvard Medical School; Biological and Biomedical Sciences Program, Harvard University; Department of Molecular and Cellular Biology, Harvard University; Department of Organismic and Evolutionary Biology, Harvard University; Department of Physics, Harvard University

## Abstract

Deep mutational scans have measured the effects of many mutations on many different proteins. Here we use a collection of such scans to perform a statistical meta-analysis of the effects of single amino acid substitutions. Specifically, we model the relative deleteriousness of each substitution in each deep mutational scan with respect to the identities of the wildtype and mutant residues, and the wildtype residue’s surface accessibility. This model explains much of the variance in mutational effects and quantifies physicochemical trends underlying them, including by yielding an empirical amino acid exchangeability matrix.

---

It is of fundamental interest in medicine [1], molecular evolution [2], and protein engineering [3] to understand the effects of mutations on proteins. However, proteins are diverse [4] and they and their properties are biophysically complex [5]. Accordingly, experimental techniques, most prominently deep mutational scanning [6], have been developed to measure the effects of mutations at high throughput on proteins of interest. Different protein properties can be assayed in this way, common effects measured being those on expression [7], binding affinity to a ligand [8], and catalytic activity [7, 9].

Machine learning models have been developed to predict the deleteriousness of mutations by comparing the mutated sequence to natural sequences that have been recorded in large repositories [16]. In rendering their predictions, these models have traditionally required input of sequences homologous to the protein of interest [12, 17, 18]. Protein language models [19, 20], recent alternatives, instead learn from a large and general corpus of protein sequences before tailoring their predictions to the given protein. Predictions of machine learning models are typically compared against deep mutational scans for validation. Databases such as MaveDB [21] and ProteinGym [15] are useful to this end because they compile deep mutational scans in a standard format. ProteinGym is designed for benchmarking models and thus also provides the predictions of several dozen machine learning models alongside experimentally measured effects.

Even since before deep mutational scanning and despite the biophysical idiosyncrasies of protein function, trends in mutational effects have been noted. Work with individual proteins revealed structural considerations in substitution tolerance [22]. It is well-recognized for example that polar substitutions to buried residues tend to be disruptive [23, 24], more so than substitutions to solvent-accessible residues [25, 26]. Similarly, substitutions to loops and turns are typically better tolerated than those to alpha helices and beta sheets [26], with mutations to proline posing a particular danger to these structured regions [27]. Indeed, methods of predicting mutational effects with respect to human disease have often incorporated physicochemical features [28–31], typically as part of a collection of information including sequence alignments.

Deep mutational scanning data now enables us to understand these physicochemical trends more precisely and with greater statistical power, an opportunity others have taken to deliver insightful analyses [32, 33]. With this data Gray et al. for example argued histidine and asparagine substitutions are the ones most demonstrative of the effects of other substitutions [34] as opposed to alanine scanning [35], and Dunham and Beltrao analyzed dozens of deep mutational scans, categorizing protein sites by patterns of mutational effects and interpreting these categories with respect to local structural context [36]. Mutational effects have been modeled with empirical amino acid exchangeability matrices [37, 38], and with the surface accessibility of the substituted residue [39, 40]. Schulze and Lindorff-Larsen combined these features to infer a separate exchangeability matrix for buried and exposed residues [41] in analyzing effects of mutations on the protein’s cellular abundance.

In this study, we measure aggregate physicochemical commonalities among mutational effects and quantify the degree to which they explain the large and varied collection of deep mutational scanning data now at our disposal. We use a statistical rank model to analyze these various deep mutational scans in concert, first with respect only to aggregate amino acid exchangeability, and then with respect also to surface accessibility. We analyze these physicochemical inferences and show that this model, on par with sequence-based machine learning models, explains much of the variance in the experimentally measured effects of single substitutions, and does so in terms of only an amino acid exchangeability matrix and a parameter for the effect of surface accessibility.

## Results

We use the deep mutational scans that have been compiled in ProteinGym, which is composed of a variety of more than 200 assays (Figure S1) and nearly 700,000 single amino acid substitutions, which are well-sampled at the level of sites; for most sites in the data, the effects of substituting the wildtype with each of the 19 other proteinogenic amino acids have been measured (Figure S2A). Although effect scores are ordered by deleteriousness within scans, due to differences in methodology they can follow widely varying scales between scans [36, 43]. Previous methods have accounted for this problem by restricting scans to those that use a specific experimental technique [41] or applying a transformation to scores toward a standard scale [36]. Deep mutational scanning scores are however consistently ordered by deleteriousness, a property which has been exploited by methods that normalize the ranks of scores within scans [37]. To better account for potential differences in the kinds of sites represented between scans, and therefore inconsistencies even in such normalized scores, we explicitly model the ranks of the effects by deleteriousness within each scan, conditioning on features of the site.

Our models are effectively Plackett-Luce models [44, 45], in that each substitution has a corresponding deleteriousness score which we infer, and in that the probability of an ordering of substitutions (by decreasing experimentally measured deleteriousness) with corresponding scores *θ*_1_, *θ*_2_, …, *θ*_*n*_ is 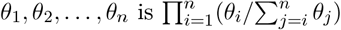. The latent deleteriousness scores are on a consistent scale: from zero to one, the latter representing maximal deleteriousness. (Although beneficial effects can be detected by deep mutational scans, we assume the best effect that can be expected is total neutrality.) In contrast to the canonical Plackett-Luce model, our models infer the *θ* by regression, analyzing the rank of each substitution in each deep mutational scan with respect to the identities of the amino acids being exchanged, and in our full model, the wildtype residue’s surface accessibility.

We first applied a model that determines the score of a particular substitution purely with respect to the identities of the amino acids being exchanged. Let *w*_*i*_ represent the identity of the wildtype amino acid of the *i*th most deleterious substitution, and let *m*_*i*_ represent that of the mutant. Then in this model the deleteriousness score of the substitution is 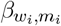 so that 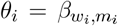 We set a prior that the set of 20 *×* 19 = 380 deleteriousness scores, for every possible exchange of amino acids, are independent and identically distributed as standard normal random variables before each being standard-logistically transformed to fall between zero and one.

Under this model, we inferred from the collected deep mutational scans the aggregate deleteriousness of substituting each amino acid for each other (Figure 1A). These scores are learned directly from substitutions in proteins that are shaped by evolution and are in at least some sense functional. For example, there are several-fold more leucine-wildtype sites in the data than cysteine-wildtype sites (Figure S2B). The inferences implicitly reflect other physicochemical trends than just the properties of the amino acids being exchanged. The relative frequencies of amino acids vary across surface accessibility (Figure S3) and secondary structure. For example, based on its tendency to disrupt secondary structure, one should expect that the typical wildtype proline is not central in alpha helices nor beta sheets, and based on their hydrophobicities, that the surface accessibility of the typical wildtype tryptophan is less than that of the typical wildtype lysine. The exchangeabilities we infer implicitly condition on these biases.

**Figure 1:**
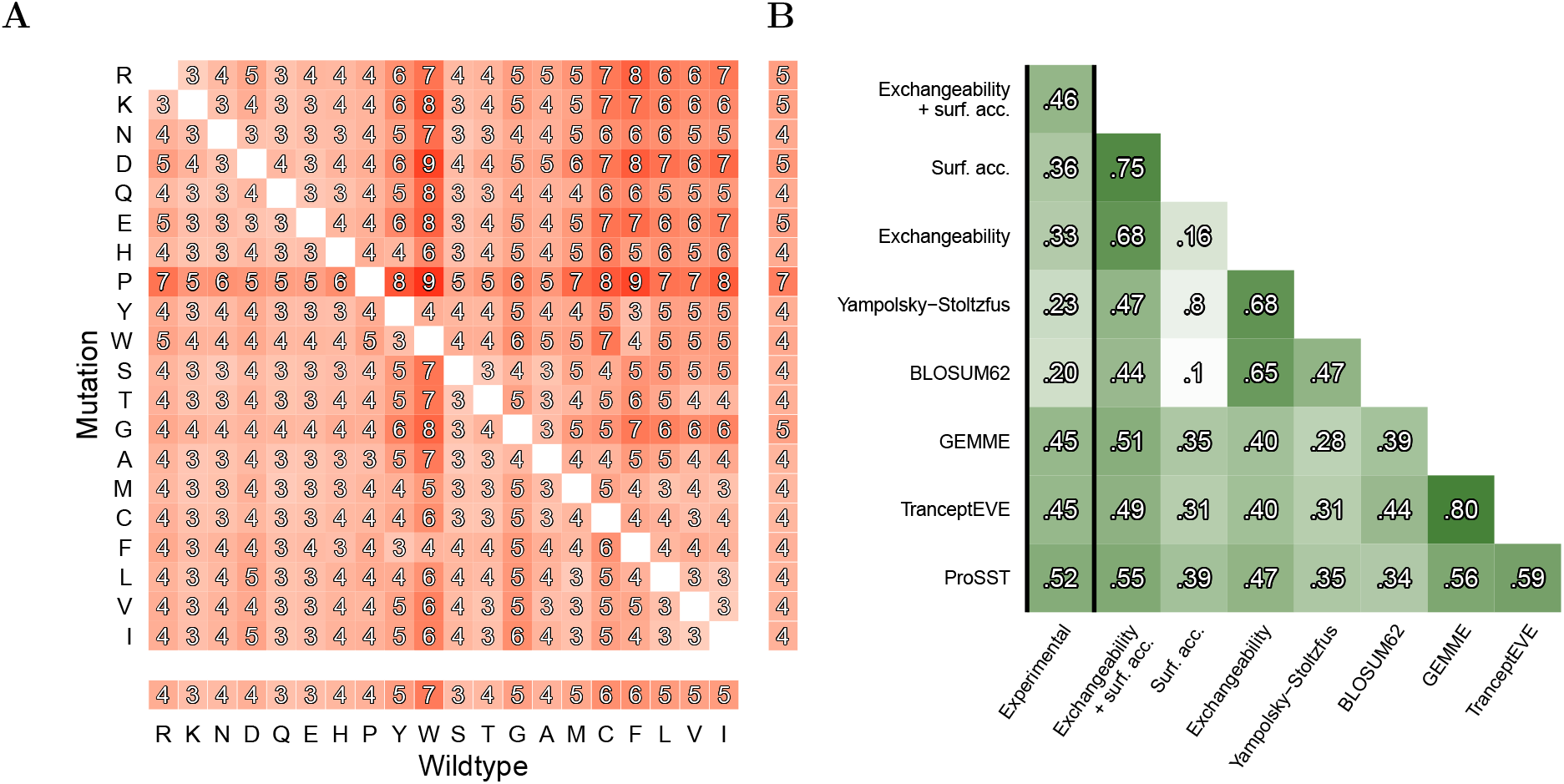
**(A)** Amino acid exchangeabilities as inferred from a model of ranks of mutational effects within deep mutational scans. Scores in the margins represent averages for the wildtype or mutant residues. Higher scores indicate greater deleteriousness, with scores rescaled *×*10 and rounded to the nearest integer for compactness. **(B)** The average per-scan Spearman rank correlation of experimental single-substitution effects, model predictions, and measures of amino acid exchangeability. Predictions of our exchangeability and combined exchangeability and surface accessibility models are from ten-fold cross-validation among proteins. Included are the Yampolsky-Stoltzfus exchangeability matrix [10] and BLOSUM62 [11]. The sequence-based machine learning models are GEMME [12], an alignment-based model, TranceptEVE [13], an alignment-based and protein language model, and ProSST [14], a structure and protein language model [15].

Altogether, these amino acid exchangeabilities explain a considerable proportion of the variance in the experimentally ranked effects (Figure 1B) and reflect known physicochemical relationships. The three least deleterious scores are assigned to the substitution of arginine for lysine followed by isoleucine for valine and valine for isoleucine. The five wildtype amino acids most prone to deleterious substitution are tryptophan followed by cysteine, phenylalanine, leucine, and isoleucine, which, excepting cysteine, are relatively large, while moderately sized amino acids are more tolerant (Figure S4). Cysteine’s obstinacy as a wildtype may be due to its unique ability to form disulfide bonds and its potential to facilitate catalytic interactions.

There are prominent asymmetries in exchangeability, in that a substitution can be more deleterious in one direction than the reverse (Figure S5) [37]. The mutant amino acid that is on average the most deleterious is proline, but proline-wildtype sites are not particularly intolerant. This contrast reflects proline’s tendency to disrupt secondary structure, while proline-wildtype sites have already in a sense accepted this biophysical limitation, just as a mutant cysteine residue would not be expected to form a disulfide bridge. The primary asymmetry is that substituting uncharged amino acids with charged ones is generally particularly deleterious, while substituting charged amino acids with uncharged ones is less so, possibly reflecting that uncharged amino acids, which are often buried in the hydrophobic core, are particularly intolerant to hydrophilic substitutes.

ProteinGym categorizes scans into broad types of protein properties that they assess: catalytic or biochemical activity, binding affinity to a target, expression, organismal growth rate, and stability. We inferred exchangeabilities from training on each of these categories separately and observed that they are fairly robust to the types of assays used to train them, with some deviations (Figure S6). Notably, replacing hydrophilic amino acids with hydrophobic ones tends to be more deleterious than usual among binding assays.

We next applied a separate model that includes surface accessibility in addition to amino acid exchangeability. In this model, the deleteriousness score of each substitution remains between zero and one, but it is computed as a logistic regression on the identities of the wildtype and mutant amino acids as well as the wildtype surface accessibility. The empirical trends of deleteriousness with respect to surface accessibility (Figure S3) indicate that this relationship is fairly consistent across substitutions, so we model it with a single parameter *β*_SA_. Specifically, we set each score *θ*_*i*_ = logistic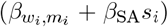, where *s*_*i*_ is the AlphaFold2-predicted wildtype surface accessibility at the site of the *i*th most deleterious substitution. We set an independent standard normal prior for each 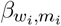 and for *β*_SA_. The inferred model is shown in Figure 2A. The inferences align fairly well with the empirical trends, and the exchangeability ranks, inde-pendent of surface accessibility, are very similar (with Spearman correlation 0.91) to those inferred by the purely amino acid-based model. Figure 2B illustrates an example of the model’s predictions.

**Figure 2:**
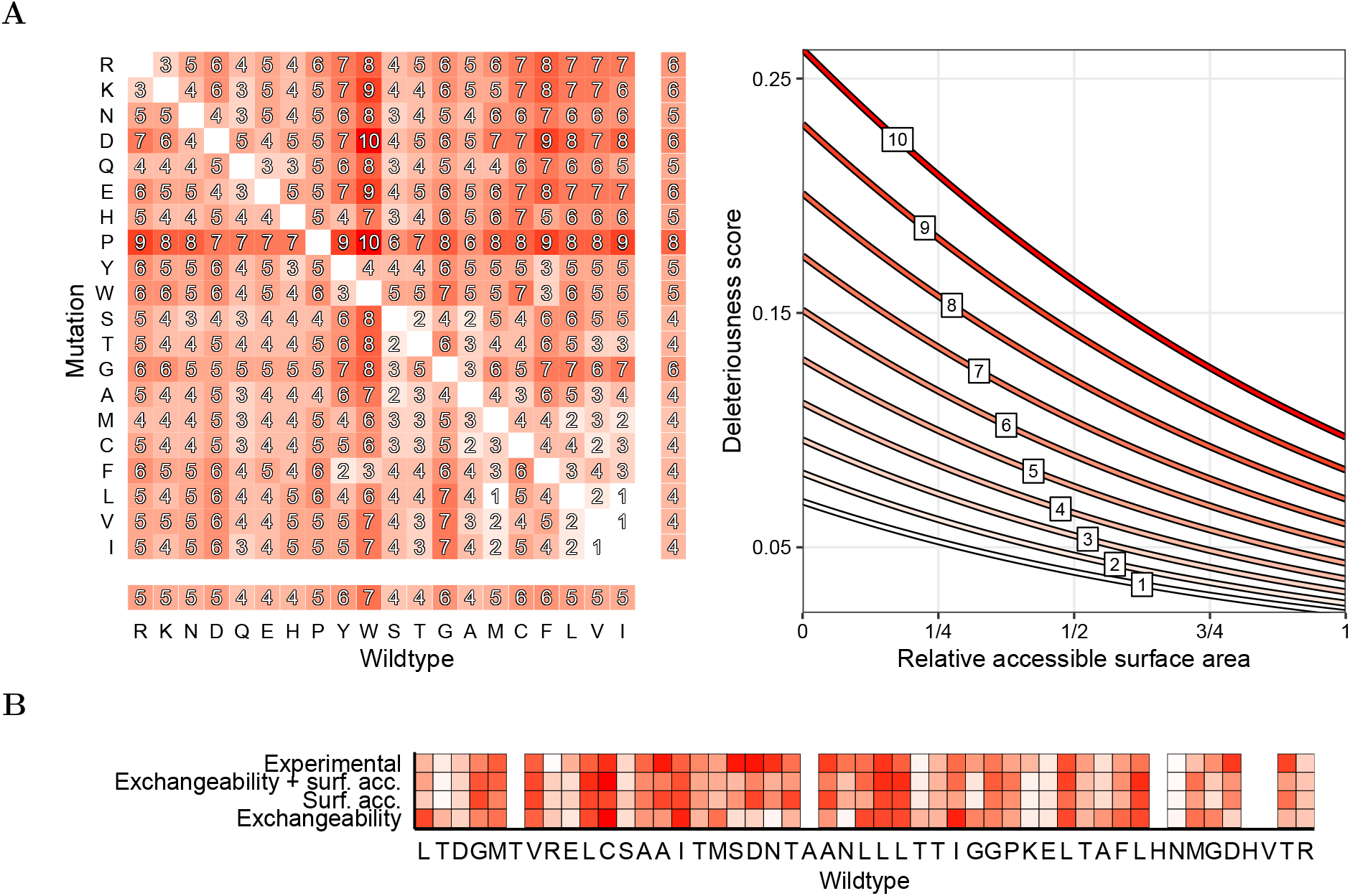
**(A)** Deleteriousness scores with respect to the amino acids being exchanged and the wildtype residue’s surface accessibility. For illustration, amino acid scores are binned from one through ten, and the deleteriousness curve with respect to surface accessibility is shown using the central score of each bin. **(B)** A demonstration of predicting the ranked deleteriousness of histidine substitutions at residues 111 to 159 of beta-lactamase based on amino acid exchangeability, surface accessibility, and both, compared to the measurements of a deep mutational scan by Firnberg et al. [42]. Some loci lack experimental measurements. The Spearman correlation of the full model’s predictions with the experiment is 0.68 in this region.

Surface accessibility alone explains the measured ranks better than exchangeability alone, and including both considerably increases the variance explained to a degree on par with sequence-based machine learning models (Figure 1B). (We also computed the average per-scan Spearman correlation of MIF [46], an inverse folding model, with deep mutation scans as 0.41, which is comparable to the others.) There is generally some but not overwhelming agreement about predictions among our models and the sequence-based models; the Spearman correlation between the predictions of our exchangeability matrix and those of TranceptEVE, for example, is 0.40. The Spearman correlation of our exchangeability and surface accessibility model with experimental measurements varies fairly widely across deep mutational scans but is more consistent among larger scans, for which the Spearman correlation ranges between approximately 0.3 and 0.6. The model is fairly similar in its average per-scan Spearman correlation across types of scans: activity (0.43), binding (0.39), expression (0.44), organismal fitness (0.39), and stability (0.57) (Figure S7).

## Discussion

By building statistical models of deep mutational scans, we collectively analyzed the relative deleteriousness of single amino acid substitutions aggregated across different proteins and protein properties with respect to amino acid exchangeability and surface accessibility. We found that these features together, on par with the predictions of even sequence-based machine learning models that involve many more parameters, tend to explain much of the variance in deleteriousness ranks within deep mutational scans.

Models that combine site-wise evolutionary conservation with amino acid exchangeability [37], and site-wise conservation with solvent accessibility [40] are also relatively simple and explain a good deal of mutational effects across a variety of types of assays. Unlike these models, ours do not condition on homologous sequence data and may overlook, for example, surface-accessible functional sites. Instead, our model essentially provides a prior over mutational effects given aggregate patterns, one we find is strong and which we anticipate will be tangibly useful toward vari-ous ends, perhaps as a prior for parameters of machine learning models especially when homologous sequence data is limited, as a baseline [38] for models that issue predictions more strongly tailored to the protein of interest, and for design, such as when a variety of still-functional variants are desired as starting points in engineering a protein. Empirical departures from the prior could also be used as evidence of functionally significant residues [41, 47].

We compared our models’ scores to the predictions of sequence-based models to understand how well our models explain the relative deleteriousness of single amino acid substitutions. These results should not be taken to mean that our aggregate physicochemical models are overall superior to sequence-based models: sequence-based models can perform tasks, including structure prediction [48] and sequence generation [20, 49], of which our models are incapable; they are unsupervised and trained on orthogonal data while ours were trained directly on deep mutational scans; and our models analyze effects of only single substitutions and therefore cannot capture epistasis, which has been observed to pervade genotype-phenotype relationships in proteins [50, 51] and which sequence models are often designed to capture [12, 18].

An immediate product of our statistical analysis was an empirical amino acid exchangeability matrix which quantifies the relative deleteriousness of each of the 380 possible amino acid substitutions. The matrix is asymmetric, in that the deleteriousness score depends not only on which amino acids are being swapped, but on which is the wildtype and which the mutant. It therefore implicitly takes into account typical physicochemical features and roles of amino acids. This matrix contrasts with substitution matrices based on evolutionary conservation, e.g. BLOSUM [11] and PAM [52], and physicochemical distance matrices [53–55], which are symmetric and do not consider mutational effects directly; substitution matrices implicitly reflect factors like codon bias and the structure of the genetic code [10] and are not intended for mutation effect prediction [17] while physicochemical distance matrices represent differences between amino acids but not directly with respect to their roles in proteins.

Yampolsky and Stoltzfus published an empirical exchangeability matrix in 2005 using data from systematic exchange studies [10], precursors to deep mutational scans. We can now draw from several times more proteins and substitutions [37]. Our matrix has a 0.64 Spearman correlation with the Yampolsky-Stoltzfus matrix and yields somewhat similar effect predictions as both this matrix and BLOSUM62 while ultimately explaining more variance in the empirical ranks (Figure 1B). We recover similar patterns in exchangeability as recent studies: our matrix has a Spearman correlation of 0.94 with one by Munro and Singh [37] and one by Høie et al. [38].

Our physicochemical models explain a substantial proportion of the variance in the relative deleteriousness of mutations without accounting for the protein properties being assayed, and the inferences from training on assay types separately are similar to those learned from the whole data. Indeed, we find that not only do surface accessibility and measures of amino acid exchangeability explain much of the variance in effects on expression [41] and stability [56], they also do well in explaining effects on other aspects of protein function. And although almost all (64 of 66) of the stability scans are relatively small and come from a single study [57] that introduced a technique for scanning domains at scale (the next-most prolific study is associated with only four scans), they are the best-explained by exchangeability and surface accessibility. These results are consistent with there being intrinsic properties of proteins, including stability [3, 32] but also perhaps collateral cellular effects such as those on gene expression and pleiotropic interactions [33, 58], that underlie assay measurements, and therefore that substitutions might, in impacting these properties, have comparable effects on higher-order properties such as binding affinity and catalytic activity.

We have shown that aggregate amino acid exchange-ability and a surface accessibility parameter explain much of the effects of single substitutions across various proteins and types of deep mutational scans. Other physicochemical features could be incorporated, although there are natural limits on how much variance in deep mutational scans can be explained given sequencing [59] and experimental noise, which can lead to divergent effect scores even between experimental replicates [60]. Idiosyncratic interactions between sites [51], the intricacies of protein function, and multi-site substitutions [61] add to the complexity of intuiting the effects of mutations. However, physicochemical trends such as those we have analyzed support that strong prior understanding of effects at short mutational distance can be formed, and may aid in the challenge to understand the grammar of proteins.

## Methods

We extracted surface accessibility data by using the software DSSP [62, 63] on the AlphaFold2-predicted [64] structures of each protein as provided by ProteinGym [15] version 1.2. We also used the metadata, model predictions, and experimental measurements for 217 deep mutational scans from this version of ProteinGym. We performed maximum a posteriori estimation of model parameters using Stan [65]. Amino acid volumes are the consensus values calculated by Perkins [66], and amino acids are ordered in figures according to Kyte and Doolittle’s hydropathy index [67]. Code can be accessed at https://github.com/berkalpay/metamutation.

## Acknowledgements

Thank you to Pascal Notin for useful discussions about ProteinGym, and to Wendy Valencia-Montoya for her feedback on the text. B.A.A. was supported by an NSF Graduate Research Fellowship under Grant No. DGE-2140743.

