## Supplementary Material for "Amino acid exchangeability and surface accessibility underpin the effects of single substitutions"

Berk A. Alpay[1], Piyush Nanda[2,3], Emma Nagy[3], Michael M. Desai[4,5]

[1]Department of Systems Biology, Harvard Medical School
[2]Biological and Biomedical Sciences Program, Harvard University
[3]Department of Molecular and Cellular Biology, Harvard University
[4]Department of Organismic and Evolutionary Biology, Harvard University
[5]Department of Physics, Harvard University

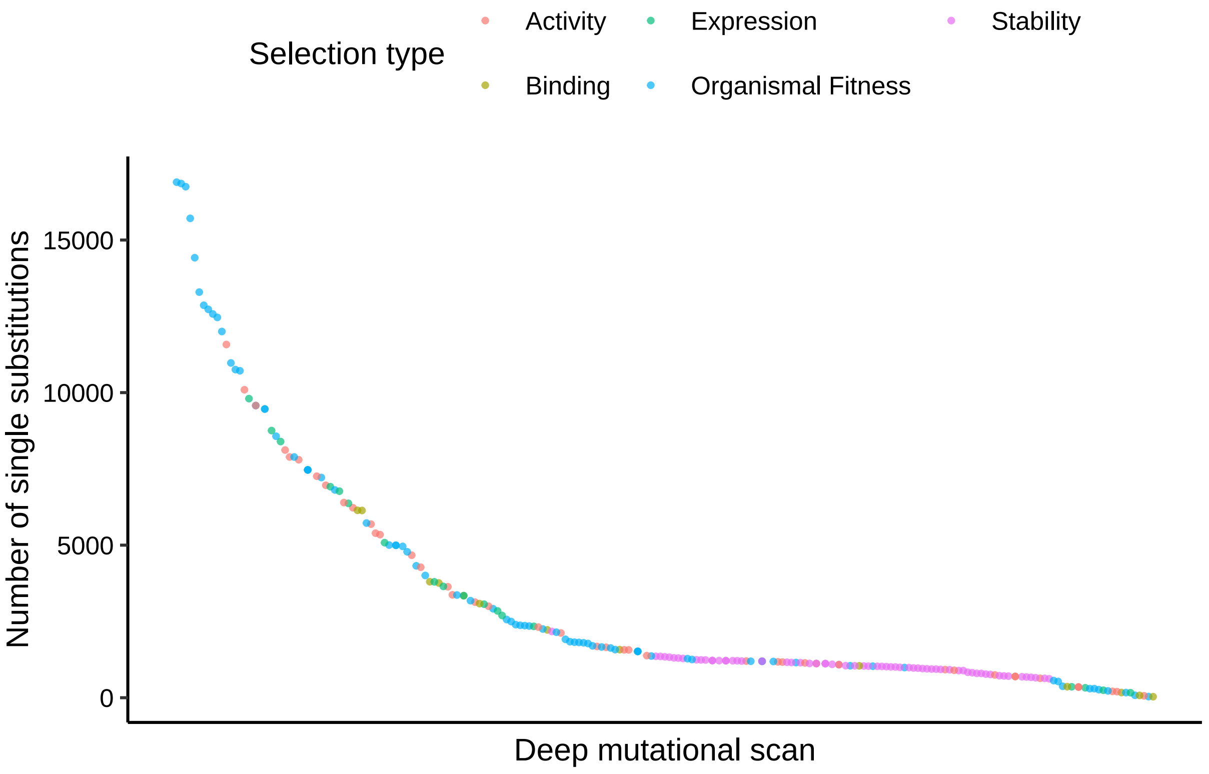


**Figure S1:** For each deep mutational scan, the selection type and the number of single amino acid substitutions measured.

**A**

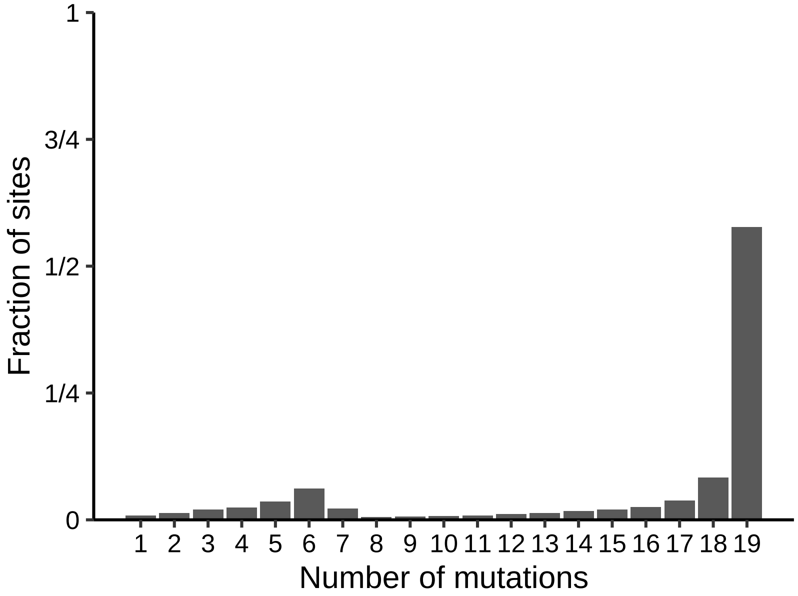


**B**

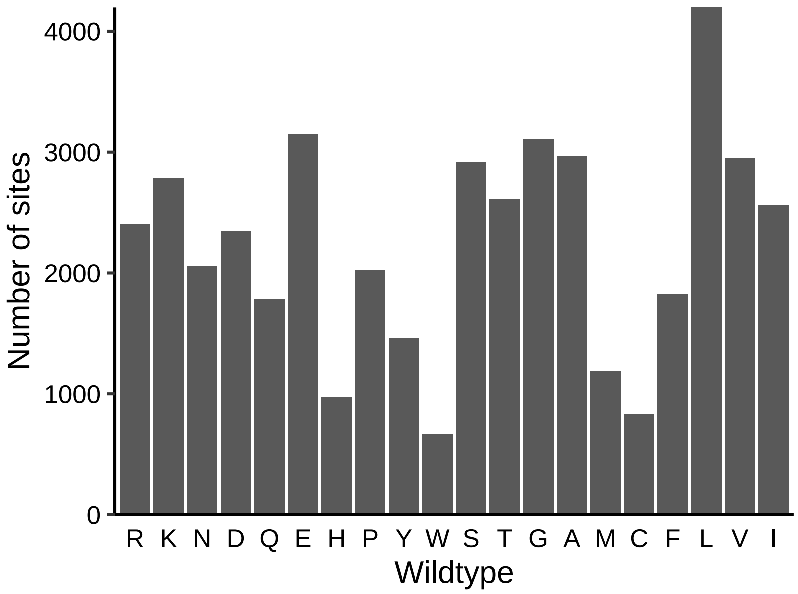


**Figure S2:** In the combined deep mutational scanning data, **(A)** the number of alternative amino acids substituted at each site, and **(B)** the number of sites at which each amino acid is the wildtype.

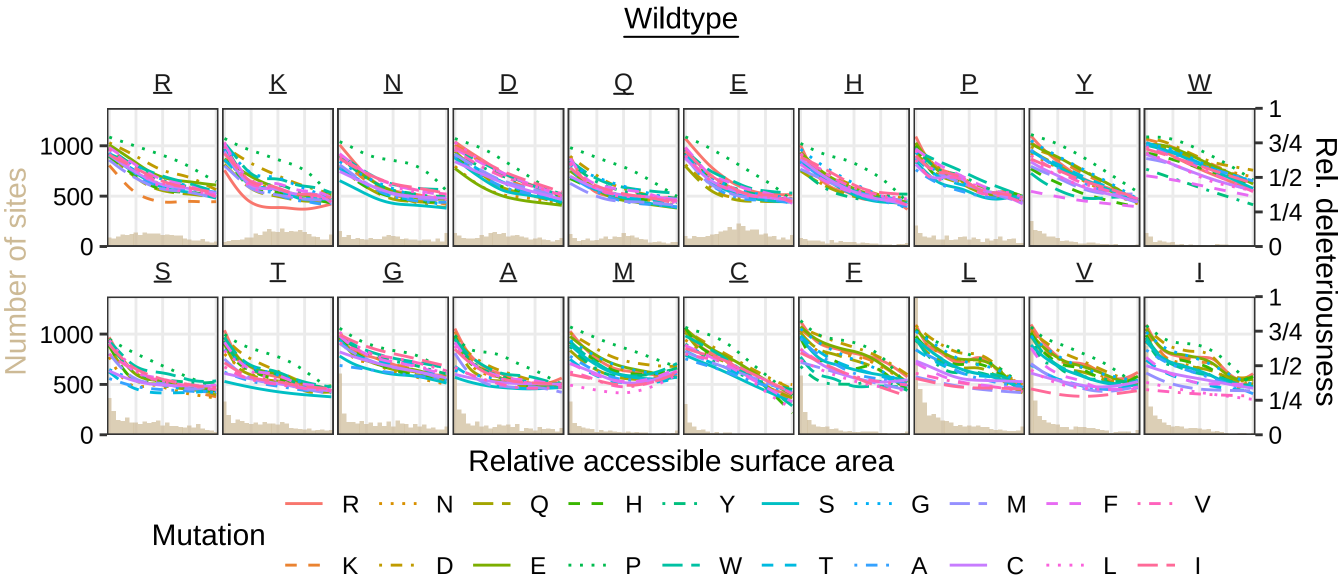


**Figure S3:** Trends in the relative deleteriousness of substitutions across deep mutational scans, with respect to the amino acids being substituted and the wildtype residue's surface accessibility. Each curve is from a separate generalized additive model, and a relative deleteriousness of 1 means the substitution is the most deleterious in its scan. Also shown is a histogram of surface accessibility for each wildtype amino acid.

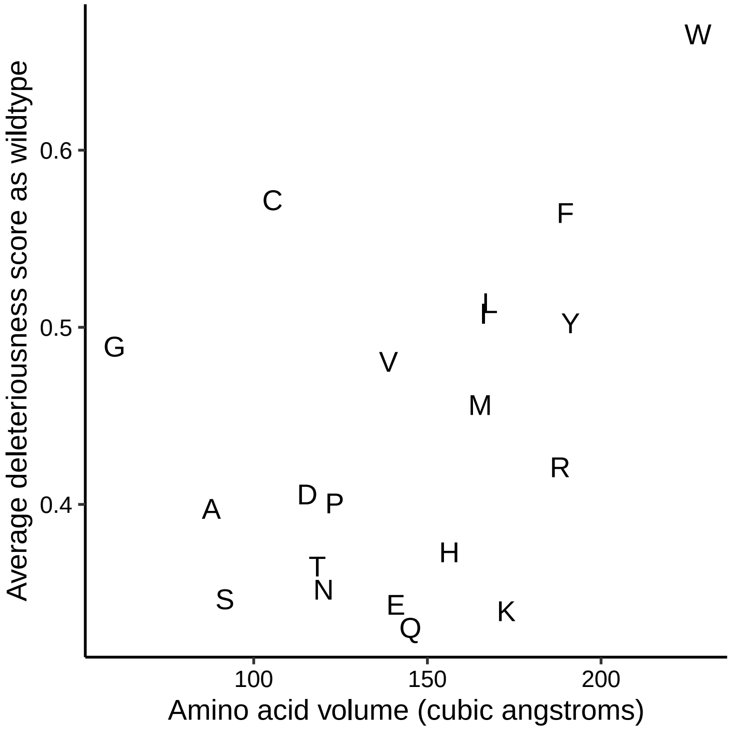


**Figure S4:** The average exchangeability of each amino acid wildtype (as in the marginal row of Figure 1A) versus its volume.

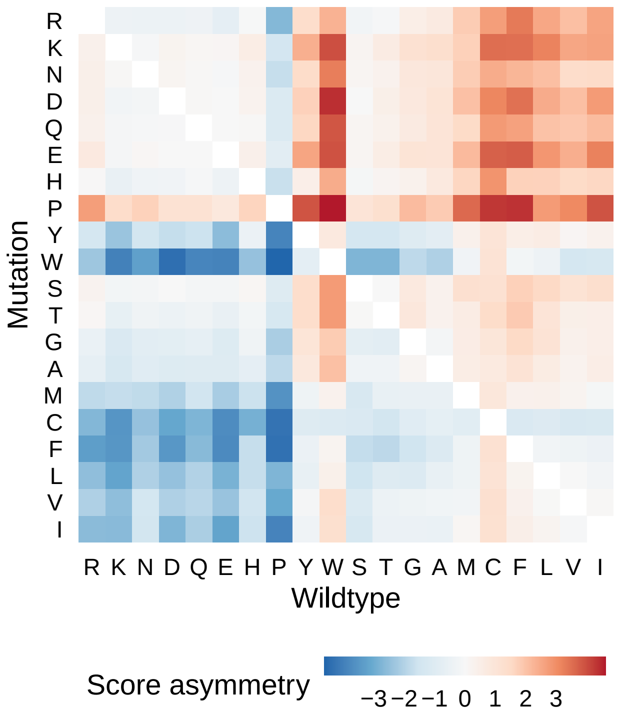


**Figure S5:** For each amino acid substitution $a$ for $b$, the deleteriousness relative to the reverse substitution $b$ for $a$, computed from Figure 1A. The greater the score asymmetry, the greater the deleteriousness of the forward substitution compared to the reverse substitution.

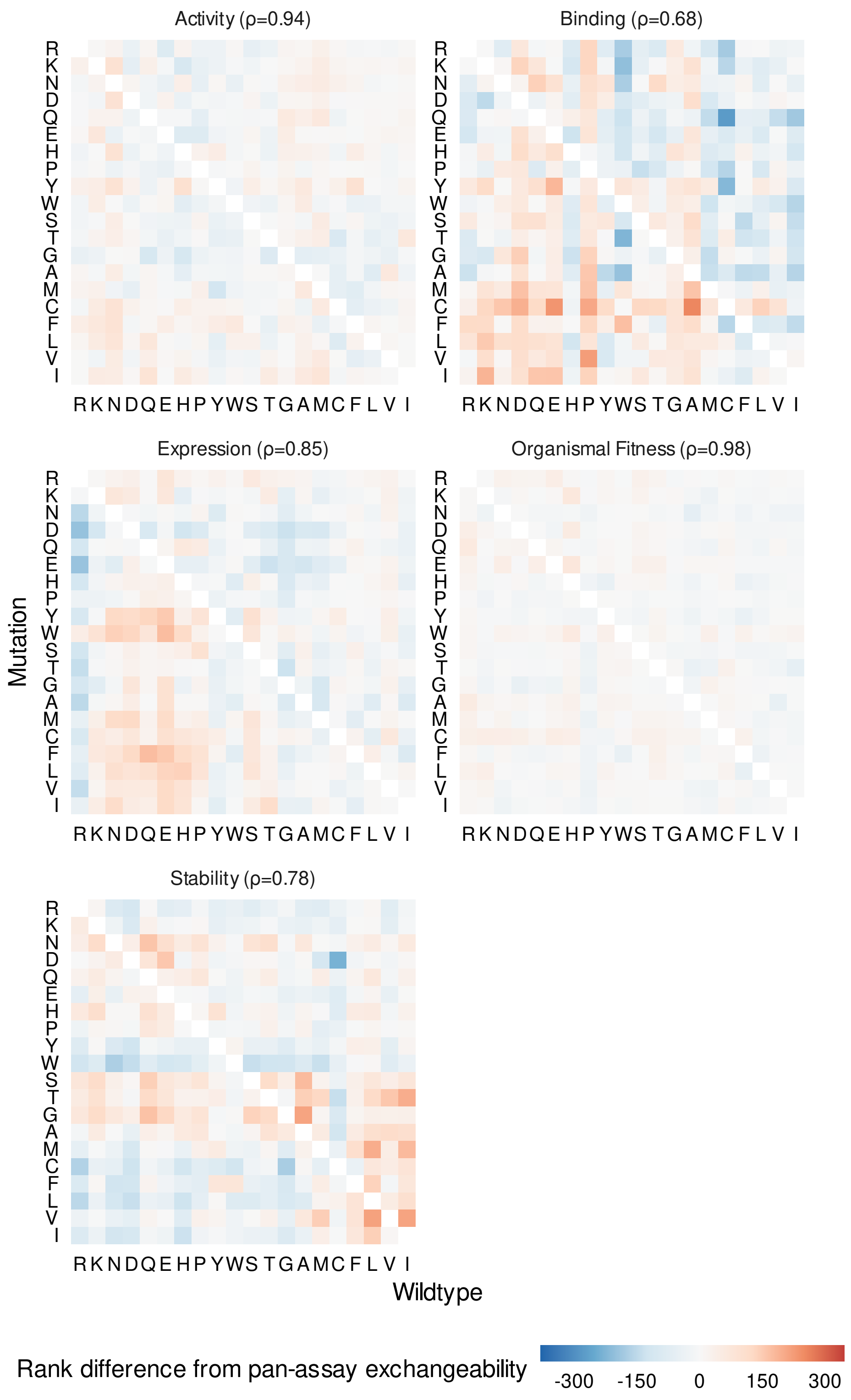


**Figure S6:** The amino acid exchangeabilities inferred by training only on assays of a certain type, shown relative to the exchangeabilities inferred from training on all types of assays (Figure 1A). Spearman correlation $\rho$ with the latter is shown for each assay type.

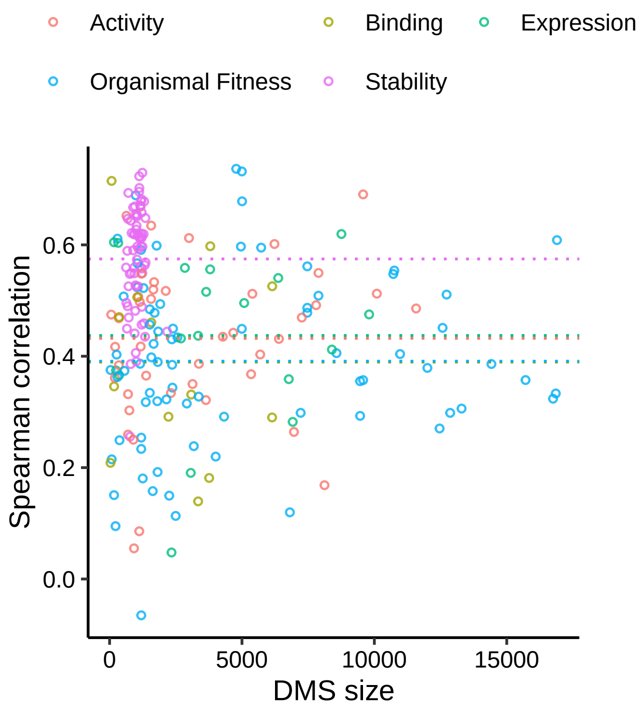


**Figure S7:** Per deep mutational scan, the Spearman correlation between experimental scores and our model of amino acid exchangeability and surface accessibility, versus the number of single-substitution effects measured. Lines represent the mean Spearman correlation for each ProteinGym-categorized type of scan.
